# *Skew*DB: A comprehensive database of GC and 10 other skews for over 28,000 chromosomes and plasmids

**DOI:** 10.1101/2021.09.09.459602

**Authors:** Bert Hubert

## Abstract

GC skew denotes the relative excess of G nucleotides over C nucleotides on the leading versus the lagging replication strand of eubacteria. While the effect is small, typically around 2.5%, it is robust and pervasive. GC skew and the analogous TA skew are a localized deviation from Chargaff’s second parity rule, which states that G and C, and T and A occur with (mostly) equal frequency even within a strand.

Most bacteria also show the analogous TA skew. Different phyla show different kinds of skew and differing relations between TA and GC skew.

This article introduces an open access database (https://skewdb.org) of GC and 10 other skews for over 28,000 chromosomes and plasmids. Further details like codon bias, strand bias, strand lengths and taxonomic data are also included. The *Skew*DB database can be used to generate or verify hypotheses. Since the origins of both the second parity rule, as well as GC skew itself, are not yet satisfactorily explained, such a database may enhance our understanding of microbial DNA.

## Background & Summary

The phenomenon of GC skew is tantalizing because it enables a simple numerical analysis that accurately predicts the loci of both the origin and terminus of replication in most bacteria and some archaea^12^.

Bacterial DNA is typically replicated simultaneously on both strands, starting at the origin of replication^3^. Both replication forks travel in the 5’ to 3’ direction, but given that the replichores are on opposite strands, topologically they are traveling in opposite directions. This continues until the forks meet again at the terminus. This means that roughly one half of every strand is replicated in the opposite direction of the other half. The forward direction is called the leading strand. Many plasmids also replicate in this way^4^.

The excess of G over C on the leading strand is in itself only remarkable because of Chargaff’s somewhat mysterious second parity rule^5^, which states that within a DNA strand, there are nearly equal numbers of G’s and C’s, and similarly, T’s and A’s. This rule does not directly follow from the first parity rule, which is a simple statement of base pairing rules.

Depending on who is asked, Chargaff’s second parity rule is so trivial that it needs no explanation, or it requires complex mathematics and entropy considerations to explain its existence^6^.

The origins of GC skew are still being debated. The leading and lagging strands of circular bacterial chromosomes are replicated very differently; it is at least plausible that this leads to different mutational biases. In addition, there are selection biases that are theorized to be involved^7^. No single mechanism may be exclusively responsible.

This article does not attempt to explain or further mystify^8^ the second parity rule or GC skew, but it may be that the contents of the *Skew*DB can contribute to our further understanding.

The *Skew*DB also contains some hard to explain data on many chromosomes. These include highly asymmetric skew, but also very disparate strand lengths. Conversely, the *Skew*DB confirms earlier work on skews in the Firmicute phylum^9^, and also expands on these earlier findings.

GC skew has often been investigated by looking at windows of DNA of a certain size. It has been found that the choice of window size impacts the results of the analysis. The *SkewDB* is therefore based exclusively on cumulative skew^10^, which sidesteps window size issues. For example, the sequence GGGCCC has a cumulative GC skew of zero, and as we progress through the sequence, this skew takes on values 1, 2, 3, 2, 1, 0.

The result of such an analysis is shown in figure 1. The analysis software fits a linear model on the skews, where it also compensates for chromosome files sequenced in the non-canonical direction, or where the origin of replication is not at the start of the file.

**Figure 1.**
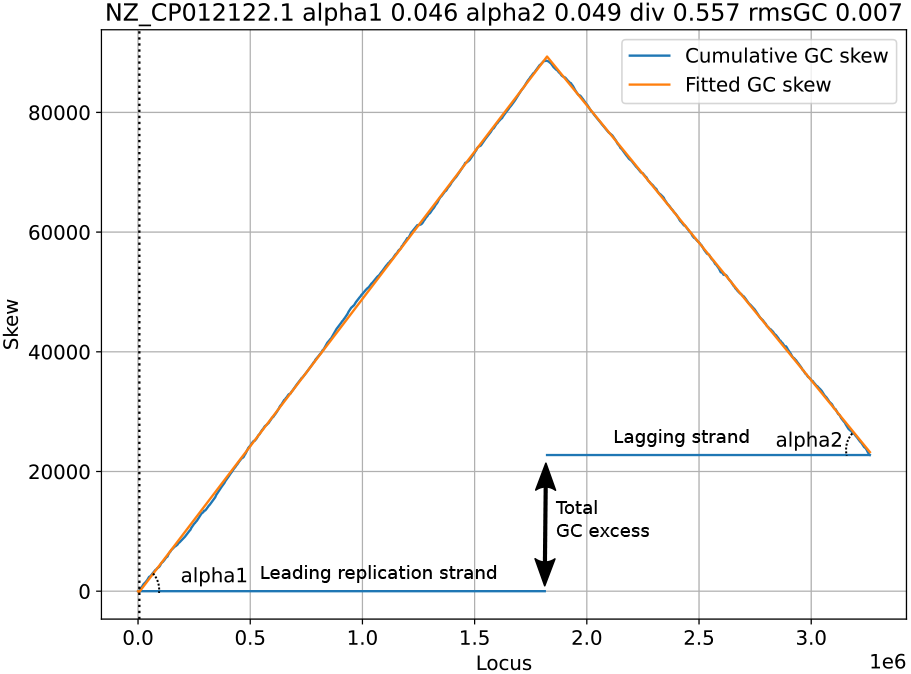
Sample graph showing *Skew*DB data for Lactiplantibacillus plantarum strain LZ95 chromosome

## Methods

The *Skew*DB analysis relies exclusively on the tens of thousands of FASTA and GFF3 files available through the NCBI download service, which covers both GenBank and RefSeq. The database includes bacteria, archaea and their plasmids.

Furthermore, to ease analysis, the NCBI Taxonomy database is sourced and merged so output data can quickly be related to (super)phyla or specific species.

No other data is used, which greatly simplifies processing. Data is read directly in the compressed format provided by NCBI. All results are emitted as standard CSV files.

In the first step of the analysis, for each organism the FASTA sequence and the GFF3 annotation file are parsed. Every chromosome in the FASTA file is traversed from beginning to end, while a running total is kept for cumulative GC and TA skew. In addition, within protein coding genes, such totals are also kept separately for these skews on the first, second and third codon position. Furthermore, separate totals are kept for regions which do not code for proteins.

In addition, to enable strand bias measurements, a cumulative count is maintained of nucleotides that are part of a positive or negative sense gene. The counter is increased for positive sense nucleotides, decreased for negative sense nucleotides, and left alone for non-genic regions. A separate counter is kept for non-genic nucleotides.

Finally, G and C nucleotides are counted, regardless of if they are part of a gene or not.

These running totals are emitted at 4096 nucleotide intervals, a resolution suitable for determining skews and shifts.

In addition, one line summaries are stored for each chromosome. These line includes the RefSeq identifier of the chromosome, the full name mentioned in the FASTA file, plus counts of A, C, G and T nucleotides. Finally five levels of taxonomic data are stored.

Chromosomes and plasmids of fewer than 100 thousand nucleotides are ignored, as these are too noisy to model faithfully. Plasmids are clearly marked in the database, enabling researchers to focus on chromosomes if so desired.

### Fitting

Once the genomes have been summarised at 4096-nucleotide resolution, the skews are fitted to a simple model.

The fits are based on four parameters, as shown in figure 1. `Alpha1` and `alpha2` denote the relative excess of G over C on the leading and lagging strands. If `alpha1` is 0.046, this means that for every 1000 nucleotides on the leading strand, the cumulative count of G excess increases by 46.

The third parameter is `div` and it describes how the chromosome is divided over leading and lagging strands. If this number is 0.557, the leading replication strand is modeled to make up 55.7% of the chromosome.

The final parameter is `shift` (the dotted vertical line), and denotes the offset of the origin of replication compared to the DNA FASTA file. This parameter has no biological meaning of itself, and is an artifact of the DNA assembly process.

The goodness-of-fit number consists of the root mean squared error of the fit, divided by the absolute mean skew. This latter correction is made to not penalize good fits for bacteria showing significant skew.

GC skew tends to be defined very strongly, and it is therefore used to pick the `div` and `shift` parameters of the DNA sequence, which are then kept as a fixed constraint for all the other skews, which might not be present as clearly.

The fitting process itself is a simplex (Amoeba) optimization over the three dimensions, seeded with the average observed skew over the whole genome, and assuming there is no shift, and that the leading and lagging strands are evenly distributed. The simplex optimization is tuned so that it takes sufficiently large steps so it can reach the optimum even if some initial assumptions are off.

For every chromosome, plasmid and skew the model parameters are stored, plus the adjusted root mean squared error.

Both for quality assurance and ease of plotting, individual CSV files are generated for each chromosome, again at 4096 nucleotide resolution, but this time containing both the actual counts of skews as well as the fitted result.

### Some sample findings

To popularize the database, an online viewer is available on https://skewdb.org/view.

#### GC and TA skews

Most bacteria show concordant GC and TA skew, with an excess of G correlating with an excess of T. This does not need to be the case however. Figure 2 is a scatterplot that shows the correlation between the skews for various major superphyla. Firmicutes (part of the Terrabacteria group) show clearly discordant skews.

**Figure 2.**
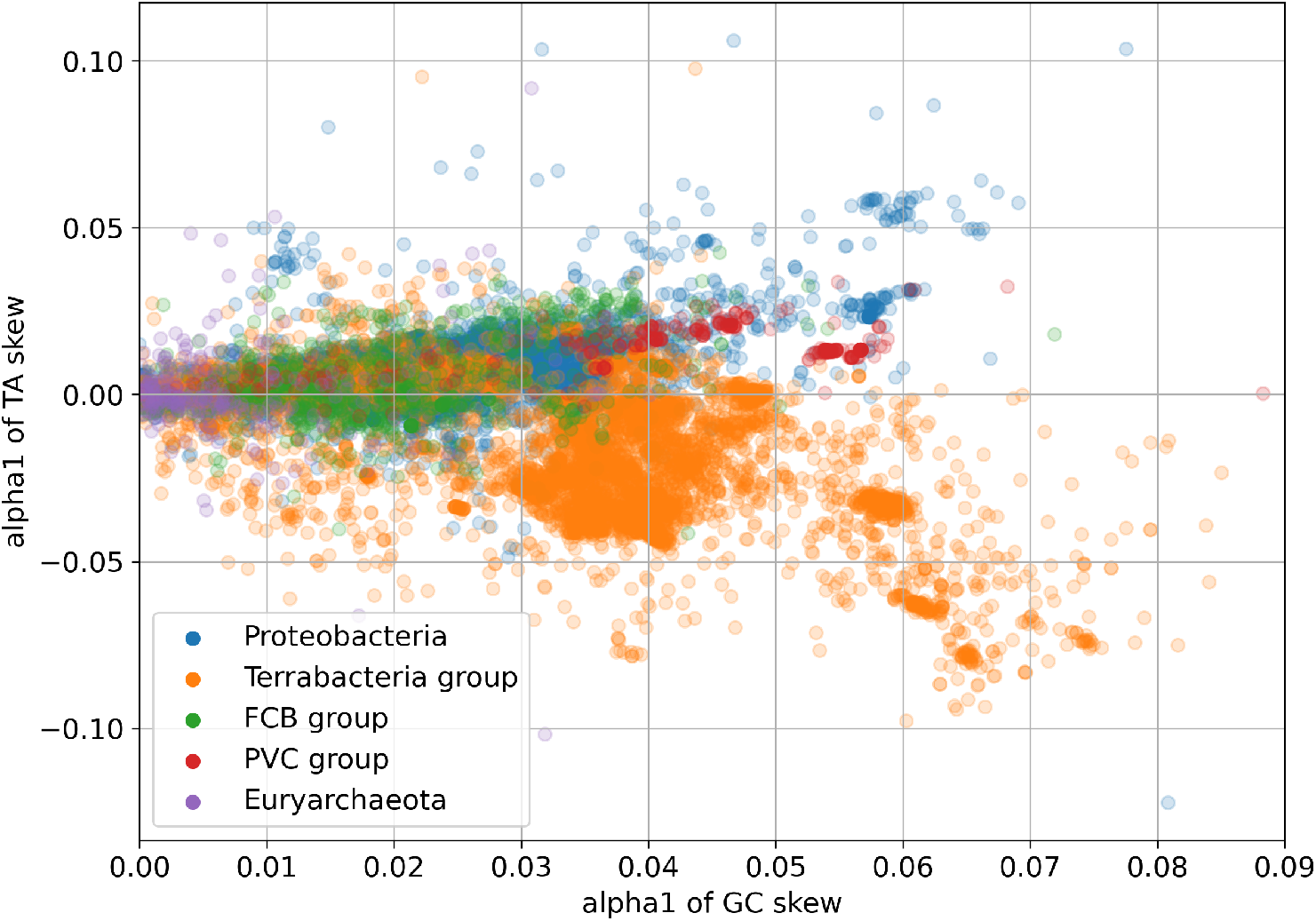
Scatter graph of 25,000 chromosomes by superphylum, GC skew versus TA skew

#### Firmicute prediction

In many bacteria, genes tend to concentrate on the leading replication strand. If the codon bias of genes is such that they are relatively rich in one nucleotide, the “strand bias” may itself give rise to GC or TA bias. Or in other words, if genes contain a lot of G’s and they huddle on the leading strand, that strand will show GC skew. As an hypothesis, we can plot the observed GC and TA skews for all Firmicutes for which we have data.

Mathematically the relation between the codon bias, strand bias and predicted GC skew turns out to be a simple multiplication. In figure 3, the x-axis represents this multiplication. The y-axis represents the GC and TA skew ratio.

**Figure 3.**
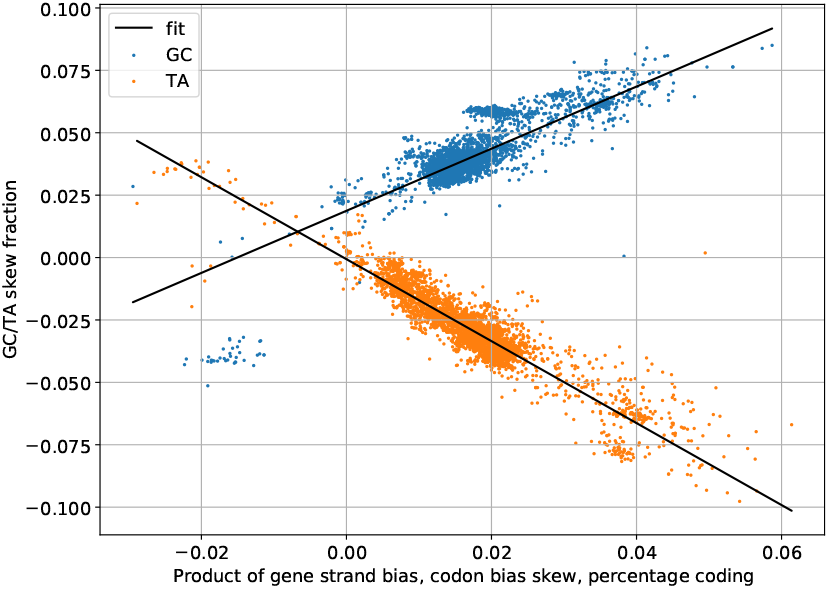
Predicted versus actual GC/TA skew for 4093 Firmicutes

It can clearly be seen that both GC and TA skew correlate strongly with the codon/strand bias product. TA skew goes to zero with the two biases, but GC skew appears to persist even in the absence of such biases.

Figure 4 shows the situation within an individual chromosome (*C. difficile*), based on overlapping 40960-nucleotide segments. On the x-axis we find the strand bias for these segments, running from entirely negative sense genes to entirely positive sense genes. The skew is meanwhile plotted on the y-axis, and here too we see that TA skew goes to zero in the absence of strand bias, while GC skew persists and clearly has an independent strand-based component.

**Figure 4.**
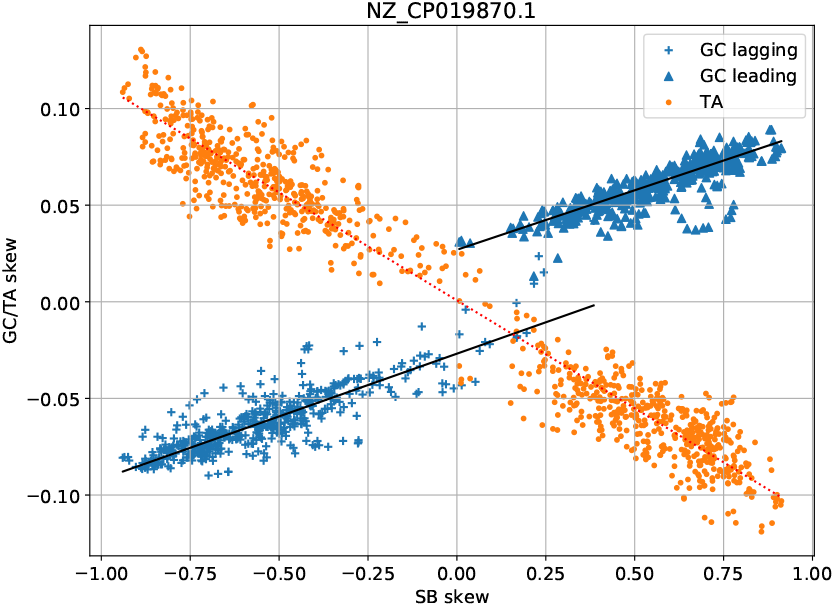
Scatter graph of codon/strand bias versus GC/TA skew for *C. difficile*

#### Asymmetric skew

The vast majority of chromosomes show similar skews on the leading and the lagging replication strands, something that makes sense given the pairing rules. There are however many chromosomes that have very asymmetric skews, with one strand sometimes showing no skew at all. In figure 5 four chromosomes are shown that exhibit such behavior. The *Skew*DB lists around 250 chromosomes where one strand has a GC skew at least 3 times bigger/smaller than the other one.

**Figure 5.**
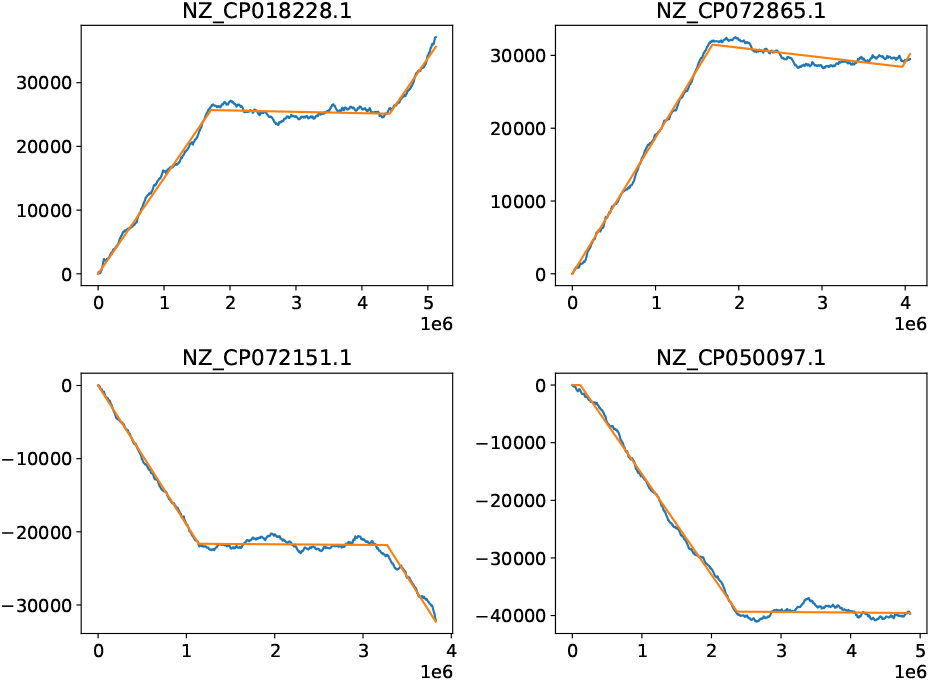
Chromosomes with asymmetric skews

#### Asymmetric strands

Bacteria tend to have very equally sized replication strands, which is also an optimum for the duration of replication. It is therefore interesting to observe that GC skew analysis finds many chromosomes where one strand is four times larger than the other strand. In figure 6 four such chromosomes are shown. The *Skew*DB lists around 100 chromosomes where one strand is at least three times as large as the other strand.

**Figure 6.**
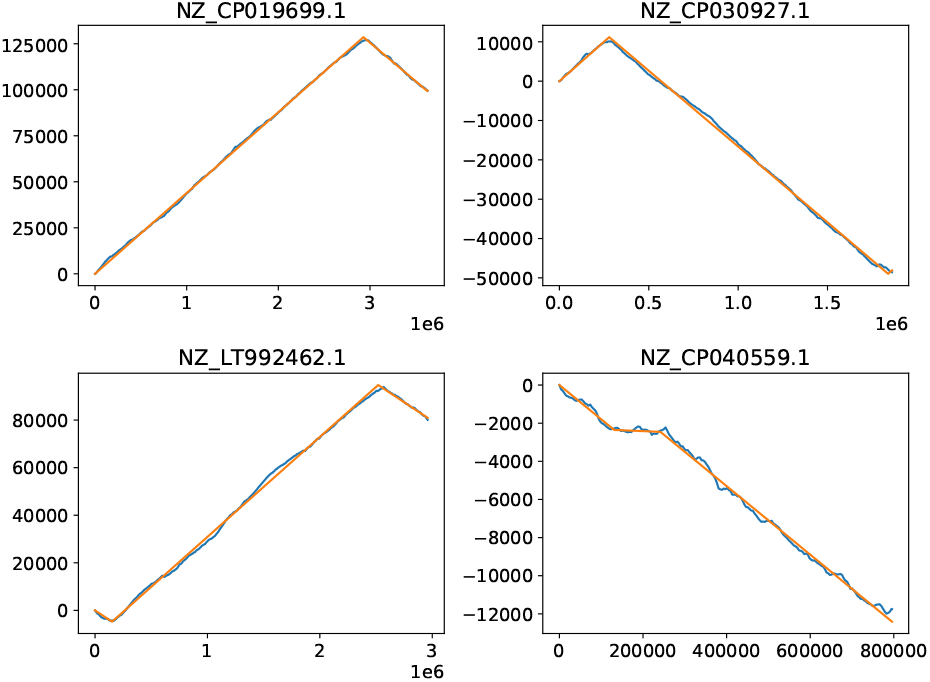
Chromosomes with differing strand lengths

#### Anomalies

In many ways, GC skew is like a forensic record of the historical developments in a chromosome. Horizontal gene transfer, inversions, integration of plasmids, excisions can all leave traces. In addition, DNA sequencing or assembly artifacts will also reliably show up in GC graphs.

Figure 7 shows GC and TA skews for *Salmonella enterica subsp. enterica serovar Concord* strain AR-0407 (NZ_CP044177.1), and many things could be going on here. The peaks might correspond to multiple origins of replication, but might also indicate inversions or DNA assembly problems.

**Figure 7.**
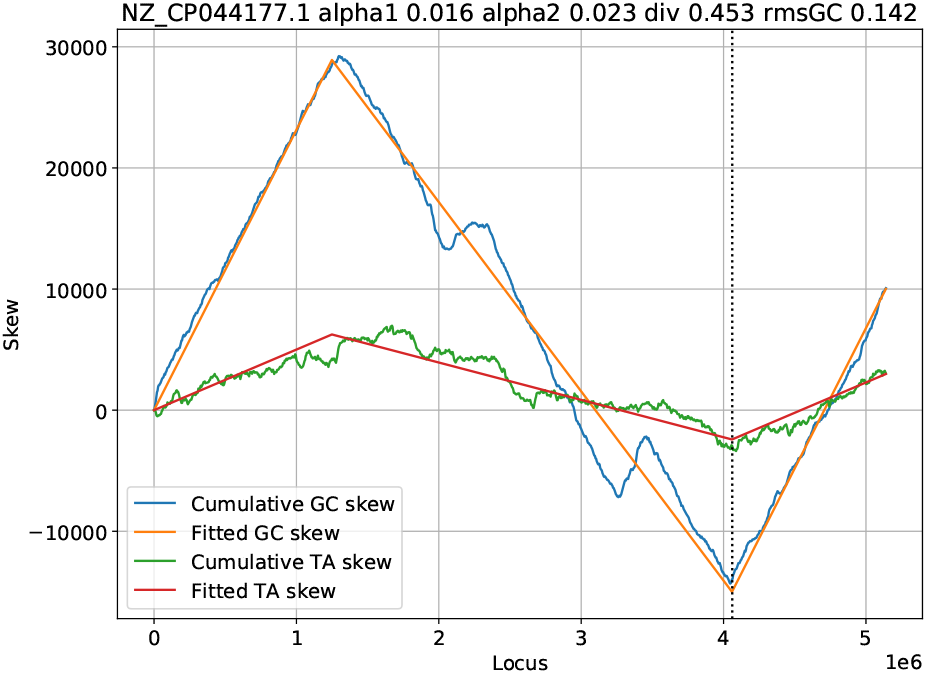
GC and TA skew for Salmonella enterica subsp. enterica serovar Concord strain AR-0407

## Data Records

The *Skew*DB consists of several CSV files: skplot.csv, results.csv, genomes.csv and codongc.csv. In addition, for each chromosome or plasmid, a separate _fit.csv file is generated, which contains data at 4096-nucleotide resolution.

skplot.csv contains all the 4096-nucleotide resolution data as one big file for all processed chromosomes and plasmids. The parameters are described in table 1.

**Table 1.**
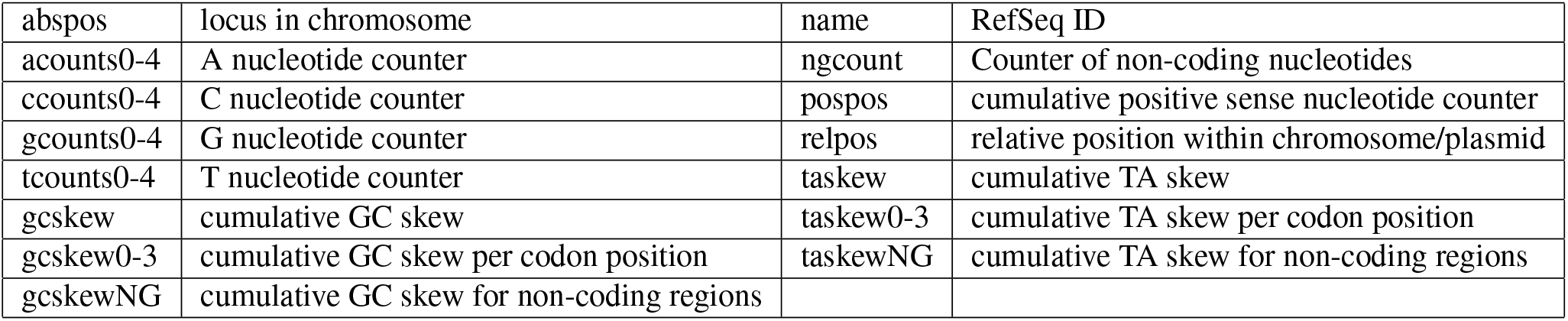
Fields of skplot.csv

results.csv meanwhile contains the details of the fits. In this table 2, all marked out squares exist. The actual fields are called alpha1gc, alpha2gc, gcRMS, alpha1ta, alpha2ta etc. DNA sequence shift and div are also specified, and they come from the GC skew. gc0-2, ta0-2 refers to codon position. gcng and tang refer to the non-coding region skews. Finally sb denotes the strand bias.

**Table 2.** Skew metrics

|  | alpha1 | alpha2 | rms | div | shift |
| --- | --- | --- | --- | --- | --- |
| gc | X | X | X | X | X |
| ta | X | X | X |  |  |
| gc0 | X | X | X |  |  |
| gc1 | X | X | X |  |  |
| gc2 | X | X | X |  |  |
| ta0 | X | X | X |  |  |
| ta1 | X | X | X |  |  |
| ta2 | X | X | X |  |  |
| gcng | X | X | X |  |  |
| tang | X | X | X |  |  |
| sb | X | X | X |  |  |

Table 3 documents the data on codon bias, also split out by leading or lagging strand found in codongc.csv.

**Table 3.** Fields in codongc.csv

|  |  |
| --- | --- |
| afrac, cfrac, gfrac, tfrac | Fraction of coding nucleotides that are A, C, G or T |
| leadafrac, leadcfrac, leadgfrac, leadtfrac | Fraction of leading strand coding nucleotides that are A, C, G or T |
| lagafrac, lagcfrac, laggfrac, lagtfrac | Fraction of lagging strand coding nucleotides that are A, C, G or T |
| ggcfrac, cgcfrac | The G and C fraction of GC coding nucleotides respectively |
| atafrac, ttafrac | The A and T fraction of AT coding nucleotides respectively |

**Table 4.**
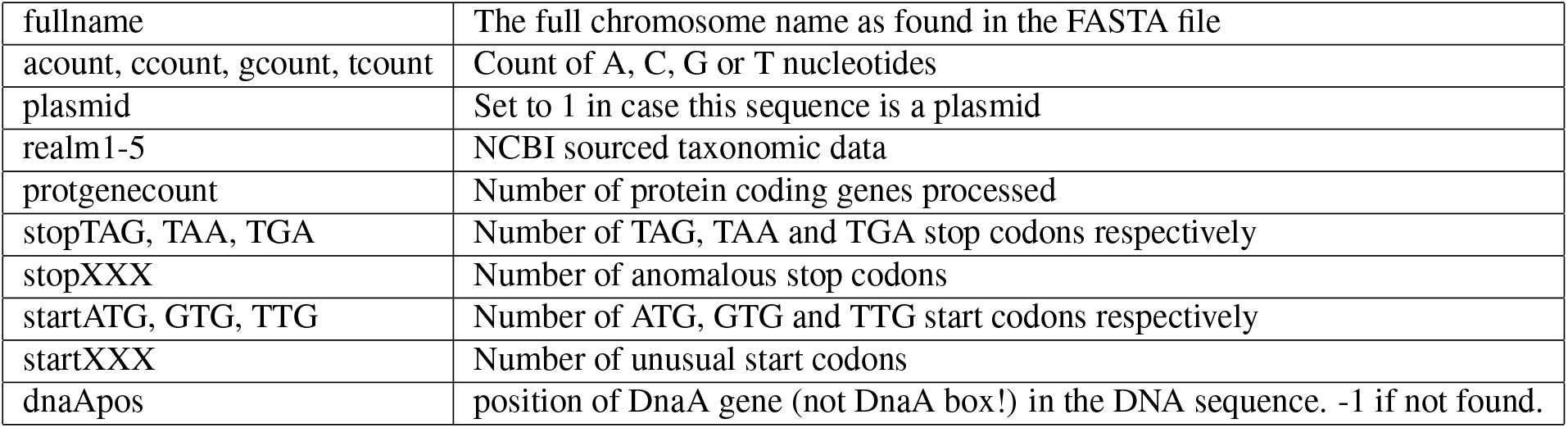
Fields in genomes.csv

The genomes.csv file contains the following fields:

Finally, the individual _fit.csv files contain fields called “Xskew” and “predXskew” to denote the observed X=gc, ta etc skew, plus the prediction based on the parameters found in results.csv.

## Technical Validation

This database models the skews of many chromosomes and plasmids. Validation consists of evaluating the goodness-of-fit compared to the directly available numbers.

The *Skew*DB fits skews to a relatively simple model of only three parameters. This prevents overfitting, and this model has proven to be robust in practice. Yet, when doing automated analysis of tens of thousands of chromosomes, mistakes will be made. Also, not all organisms show coherent GC skew.

Figure 8 shows 16 equal sized quality categories, where it is visually clear that the 88% best fits are excellent. It is therefore reasonable to filter the database on *RMS_gc_* < 0.1067. Or conversely, it could be said that above this limit interesting anomalous chromosomes can be found.

**Figure 8.**
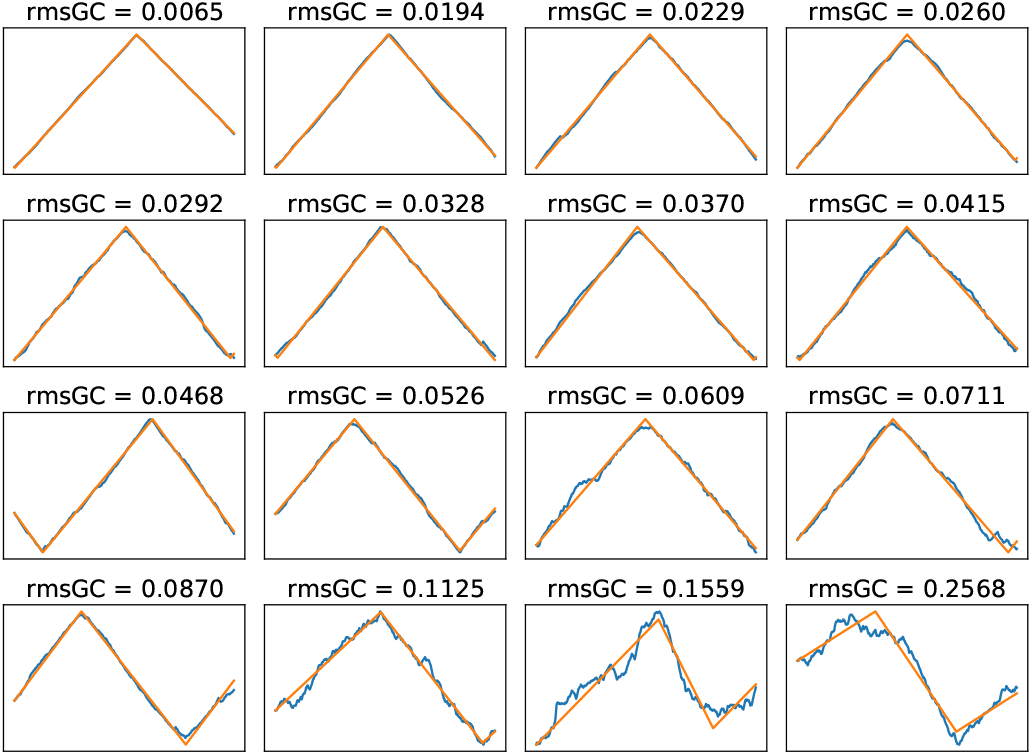
*Skew*DB fits for 16 equal sized quality categories of bacterial chromosomes

The DoriC database^2^ contains precise details of the location of the origin of replication. 2267 sequences appear both in DoriC and in the *Skew*DB. The DoriC origin of replication should roughly be matched by the “shift” metric in the *Skew*DB (but see Usage notes). For 90% of sequences appearing in both databases, there is less than 5% relative chromosome distance between these two independent metrics. This is encouraging since these two numbers do not quite measure the same thing.

On a similar note, the DnaA gene is typically (but not necessarily) located near the origin of replication. For around 80% of chromosomes, DnaA is found within 5% of the *Skew*DB “shift” metric. This too is an encouraging independent confirmation of the accuracy of the data.

Finally, during processing numbers are kept of the start and stop codons encountered on all protein coding genes on all chromosomes and plasmids. These numbers are interesting in themselves (because they correlate with GC content, for example), but they also match published frequencies, and show limited numbers of non-canonical start codons, and around 0.005% anomalous stop codons. This too confirms that the analyses are based on correct (annotation) assumptions.

## Usage Notes

The existential limitation of any database like the *Skew*DB is that it does not represent the distribution of organisms found in nature. The sequence and annotation databases are dominated by easily culturable microbes. And even within that selection, specific (model) organisms are heavily oversampled because of their scientific, economic or medical relevance.

Because of this, care should be taken to interpret numbers in a way that takes such over- and undersampling into account. This leaves enough room however for finding correlations. Some metrics are sampled so heavily that it would be a miracle if the unculturable organisms were collectively conspiring to skew the statistics away from the average. In addition, the database is a very suitable way to test or generate hypotheses, or to find anomalous organisms.

Finally it should be noted that the *Skew*DB tries to precisely measure the skew parameters, but it makes no effort to pin down the Origin of replication exactly. For such uses, please refer to the DoriC database^2^. In future work, the *Skew*DB will attempt to use OriC motifs to improve fitting of this metric.

On https://skewdb.org an explanatory Jupyter^11^ notebook can be found that uses Matplotlib^12^ and Pandas^13^ to create all the graphs from this article, and many more. In addition, this notebook reproduces all numerical claims made in this work. The *Skew*DB website also provides links to informal articles that further explain GC skew, and how it could be used for research.

## Code availability

The *Skew*DB is produced using the [Antonie DNA processing software](https://github.eu/berthubert/antonie2), which is open source. In addition, once the RefSeq and GenBank files have been created and from the NCBI website, the pipeline is fully automated and reproducible.

A [GitHub repository](https://github.com/berthubert/skewdb-articles) is available for this article, which includes this reproducible pipeline, plus a script that regenerates all the graphs and numerical claims from this paper.

## Acknowledgements

I would like to thank Bertus Beaumont for helping me to think like a biologist, and Jason Piper for regularly pointing me to the relevant literature. In addition, I am grateful that Felix Hol kindly allowed me to field test my software on his DNA sequences^14^. Twitter users @halvorz and @Suddenly_a_goat also provided valuable feedback.

## Author contributions statement

B.H. did all the work.

